# Divergent *MLS1* promoters lie on a fitness plateau for gene expression

**DOI:** 10.1101/026427

**Authors:** Andrew C. Bergen, Gerilyn M. Olsen, Justin C. Fay

**Affiliations:** Molecular Genetics and Genomics Program, Washington University, St. Louis, MO; Washington University, St. Louis, MO; Department of Genetics, Washington University, St. Louis, MO; Center for Genome Sciences and Systems Biology, Washington University, St. Louis, MO

**Keywords:** yeast, evolution, expression, *MLS1*, binding sites

## Abstract

Qualitative patterns of gene activation and repression are often conserved despite an abundance of quantitative variation in expression levels within and between species. A major challenge to interpreting patterns of expression divergence is knowing which changes in gene expression affect fitness. To characterize the fitness effects of gene expression divergence we placed orthologous promoters from eight yeast species upstream of malate synthase (*MLS1*) in *Saccharomyces cerevisiae*. As expected, we found these promoters varied in their expression level under activated and repressed conditions as well as in their dynamic response following loss of glucose repression. Despite these differences, only a single promoter driving near basal levels of expression caused a detectable loss of fitness. We conclude that the *MLS1* promoter lies on a fitness plateau whereby even large changes in gene expression can be tolerated without a substantial loss of fitness.

## INTRODUCTION

Changes in gene regulation are thought to play an important role in evolution (King and Wilson 1975; Wray 2007; Carroll 2008). While there are many examples of *cis*-regulatory changes underlying diverged phenotypes (Wray 2007; Gaunt and Paul 2012), phenotypes more often map to changes in protein coding sequences (Hoesktra and Coyne 2007; Stern and Orgogozo 2008; Martin and Orgogozo 2013; Fay 2013). One reason for the fewer number of phenotypes attributable to *cis*-regulatory mutations is the greater difficulty in demonstrating their influence on a phenotype (Stern and Orgogozo 2008). As such, most of our understanding of regulatory evolution is based on gene expression and *cis-*regulatory sequence divergence irrespective of downstream phenotypes.

One general feature of regulatory evolution that has emerged is conservation of qualitative patterns of gene expression despite divergence in the *cis-*regulatory sequences driving expression. Quantitative studies of gene expression levels have shown that there is an abundance of quantitative variation within and between species (Whitehead and Crawford 2006; Gordon and Ruvinsky 2012). Yet, qualitative patterns of activation, repression and tissue-specific expression are generally conserved across distantly related species (Gasch *et al.* 2004; Chan *et al.* 2009). In comparison, studies of *cis*-regulatory sequences have shown that gain and loss of transcription factor binding sites is common (Moses *et al.* 2006; Doniger and Fay 2007; Kim *et al*. 2009; Bradley *et al.* 2010; Schmidt, *et al*. 2010; Yokoyama *et al.* 2014), and that between distantly related species, *cis*-regulatory sequences often diverge to the extent that the sequences are unalignable (Wratten *et al* 2006; Hare *et al* 2008; Venkatarum and Fay 2010; Arnold *et al.* 2014).

The binding site turnover model explains how gene regulation can be conserved while *cis*-regulatory sequences diverge (Ludwig *et al.* 1998; 2000; Dermitzakis and Clark 2002; Dermitzakis *et al.* 2003). Under this model, gain and loss of equivalent binding sites within the same regulatory sequence enables high rates of divergence without changes in gene regulation. The binding site turnover model is supported by striking demonstrations that diverged *cis*-regulatory sequences from distantly related species drive very similar patterns of gene expression when placed in the same genome (Romano and Wray 2003; Ruvinsky and Ruvkun 2003; Markstein *et al.* 2004; Fisher *et al.* 2006; Wratten *et al.* 2006; Hare *et al.* 2008; Swanson *et al.* 2011; Arnold *et al* 2014). Over long time periods, divergence in *cis*-regulatory sequences may also be facilitated by transcriptional rewiring, whereby different binding sites can be substituted for one another (Tsong *et al.* 2006; Tuch *et al.* 2008). However, the decrease in regulatory conservation as *cis*-regulatory sequences are placed into more distantly related genomes (Gordon and Ruvinsky 2012; Barriere and Ruvinsky 2014) implies that there are limits to the compatibility of *cis*-regulatory sequences with distantly related *trans*-environments.

A major barrier to interpreting patterns of gene expression conservation and divergence is that their influence on outward phenotypes or fitness is unknown. While in some instances patterns of gene expression divergence themselves are indicative of fitness effects, in most cases expression divergence is assumed to be neutral (Fay and Wittkopp 2008). For example, there is evidence that subtle but consistent changes in the expression of genes in the same pathway or biological process influence fitness (Bullard *et al.* 2010; Fraser *et al.* 2010). A further complication is that fitness may depend not only on expression levels. The temporal or developmental patterns of expression may also influence fitness. For example, by comparing the distribution of mutation effects to naturally occurring polymorphism in the gene *TDH3*, Metzger *et al.* (2015) inferred that there is abundant purifying selection against mutations that increase cell to cell variation in expression levels. Overall, testing whether *cis*-regulatory sequences have diverged in their ability to integrate transcription factors, nucleosome positioning and core transcriptional machinery into proper expression is challenging. However, the consequences of any meaningful regulatory changes should be reflected in fitness.

The direct effects of gene expression on fitness are not often characterized. Ludwig *et al.* (2005) found complementation of diverged enhancers, though none of the transgenic constructs rescued wild type fitness levels. Using an inducible promoter, a fitness plateau was found for *LCB2* gene expression in yeast (Rest *et al.* 2013). In this case, fitness increased with gene expression levels, but above a certain level no further changes in fitness were observed (Rest *et al.* 2013). While there are also many examples of expression changes that underlie phenotypes likely to influence fitness (Hoesktra and Coyne 2007; Stern and Orgogozo 2008), it is difficult to make generalizations about the nature of these expression changes.

Here, we examine the effects of promoter divergence on both gene expression and fitness in yeast using the malate synthase (*MLS1*) promoter. As part of the glyoxylate cycle, *MLS1* is induced in the absence of fermentable carbon and repressed in the presence of glucose (Turcotte *et al*. 2010). *MLS1* converts acetyl-CoA into malate and is necessary for gluconeogenesis and growth on non-fermentable carbon sources (Hartig *et al.* 1992). Additionally, *MLS1* has a well characterized promoter, where the main transcription factor binding sites and regions necessary for activation and repression have previously been identified (Caspary *et al.* 1997). Activation of *MLS1* occurs through two Abf1 binding sites, responsible for basal expression levels, and two Cat8 binding sites, responsible for its large increase in expression following depletion of glucose (Caspary *et al.* 1997). Cat8 binding sites have also been shown to be bound by Sip4 (Roth *et al.* 2004). The main transcription factors that control *MLS1* expression are conserved across species. Activation of *MLS1* by the transcription factor *CAT8* is conserved in *Kluyveromyces lactis*, a species that split before the whole genome duplication and a shift in metabolism from respiratory to fermentative growth in the present of oxygen (Georis *et al.* 2000). Repression of *MLS1* occurs through a Mig1 site (Caspary *et al.* 1997). It has been shown that a *MIG1* gene deletion in *Saccharomyces cerevisiae* can be rescued by *MIG1* from *Candida utilis* (Delfin *et al.* 2001) and *K. lactis* (Cassart *et al.* 1995), indicating that *MIG1* has conserved its general function as well.

Assays of orthologous *cis*-regulatory sequence function in a single species background have previously been valuable in understanding how they evolve. For example, loss of function can be caused by incompatibility between *cis*-regulatory sequences and trans-acting factors (Barriere *et al.* 2012) or by gain and loss of binding sites within the same *cis*-regulatory sequence (Ludwig *et al.* 2000). Here, we place orthologous *MLS1* promoters from eight different yeast species into *Saccharomyces cerevisiae* to determine what selective constraints act on this promoter as well as what expression levels and dynamics *S. cerevisiae* requires for *MLS1* function. We expected and found that orthologous promoters caused differences in gene expression levels while maintaining the general pattern of activation and repression. We then used competitive growth assays to show that despite varying expression levels, all but one of the species’ promoters completely rescues competitive fitness in *S. cerevisiae*. Our results demonstrate that most of the diverse configurations of binding sites within the *MLS1* promoter drove expression levels that can be tolerated without substantial fitness effects. We conclude that evolution of the *MLS1* promoter supports a model of neutral or nearly neutral expression divergence above a certain threshold required for normal function.

## RESULTS

### High sequence divergence with conservation of binding site content

To characterize sequence divergence in the *MLS1* promoter we examined the noncoding sequences between *MLS1* and the codon region of the upstream gene in eight yeast species. Similar to genome-wide patterns of promoter evolution in yeast (Venkataram and Fay 2010), the *MLS1* promoter exhibits: 1) an abundance of conserved sites under purifying selection based on a substitution rate of 0.15 compared to the synonymous substitution rate of 0.21 in the *MLS1* coding region (Fay and Benavides 2005); 2) no significant alignment between *S. cerevisiae* and the more distantly related non-*Saccharomyces* species; and 3) good matches to known binding sites in most of the species’ promoter sequence (Figure 1 and Figure S1). Binding sites known to regulate *MLS1* expression in *S. cerevisiae* are two activation sites, which could be bound by either Cat8 or Sip4, a Mig1 repression site and two Abf1 sites thought to be involved in basal expression (Caspary et al 1997; Roth *et al.* 2004). Although the number, position and orientation of matching binding sites are different in all but the *Saccharomyces* species, they contain good matches to the known binding sites. The one exception is *N. castellii*, which lacks a good TATA and Mig1 site. However, the binding site scores tend to be lower in more distantly related species, as measured by the total binding affinity predicted for each promoter (Table S1).

**Figure 1.**
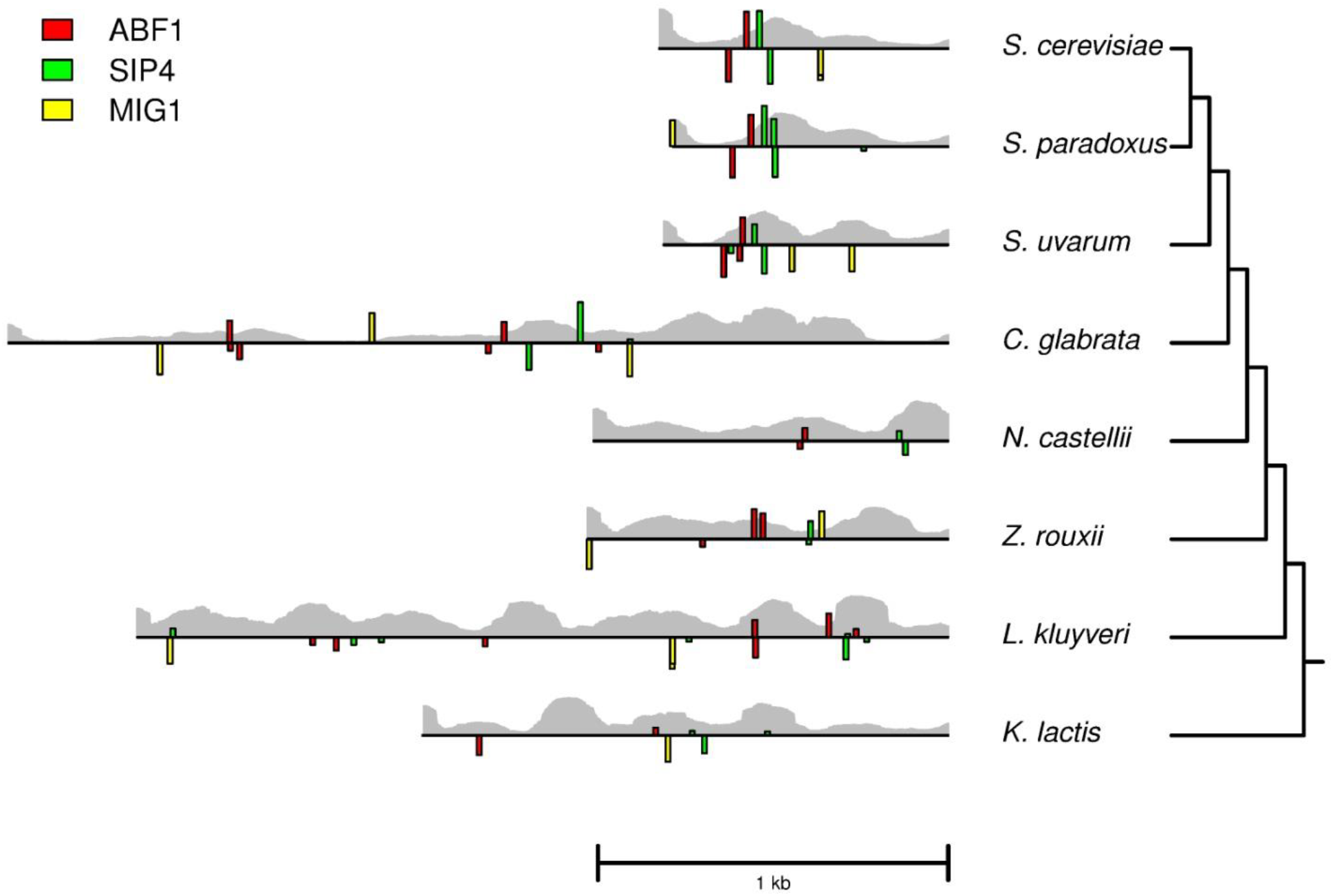
Transcription factor binding and nucleosome occupancy predictions for *MLS1* promoters. The noncoding region upstream of *MLS1* from eight yeast species, where the heights of colored bars represent the scores of predicted binding sites for Abf1, Sip4, and Mig1 based on position weight matrices. Bars above each line represent sites on the forward strand and those below represent sites on the reverse strand. The probability of nucleosome occupancy at each base pair along the promoters is represented by the height of the grey bars in the background.

To examine potential differences in nucleosome occuppancy we used a sequence-based prediction method (Kaplan *et al.* 2009), which matches in vivo measurements of nucleosome occuppancy at *MLS1* in *S. cerevisiae*. Similar to other noisy promoters, *MLS1* is characterized by a TATA element with nucleosomes positioned over its other binding sites in the presence of glucose. In ethanol, in vivo nucleosome occupancy goes down but not completely, typical of noisy gene expression found for many condition-specific genes (Kaplan *et al.* 2009). Occupancy predictions for the other yeast species are similar in that binding sites are often occupied (Figure 1).

### Conserved regulatory patterns despite changes in gene expression levels

To test whether differences in the position, orientation and slight changes in binding affinity affect gene expression, we placed each of the 8 species’ noncoding regions upstream of *S. cerevisiae MLS1*. All the promoters caused significant activation of *MLS1* in ethanol compared to glucose, ranging from 5.9 to 188-fold increase in expression (Table S2), demonstrating conservation of the response to carbon source. However, there is a general trend of a loss of both repression and activation in the most distantly related species (Figure 2A-B). Except for *K. lactis,* all non-*Saccharomyces* species’ promoters drove significantly lower expression compared to *S. cerevisiae* (Bonferroni correct p-value < 0.05, Figure 2A). In glucose, both *N. castellii* and *L. kluyveri* were not as well repressed as *S. cerevisiae* (Bonferroni corrected p-value < 0.05, Figure 2B). Interestingly, the *N. castellii* promoter does not contain either a Mig1 repressor site (Figure 1) or a proximal TATA element (Figure S1). The absence of a TATA box is known to correspond with a small dynamic range of expression (Basehoar *et al.*, 2004).

**Figure 2.**
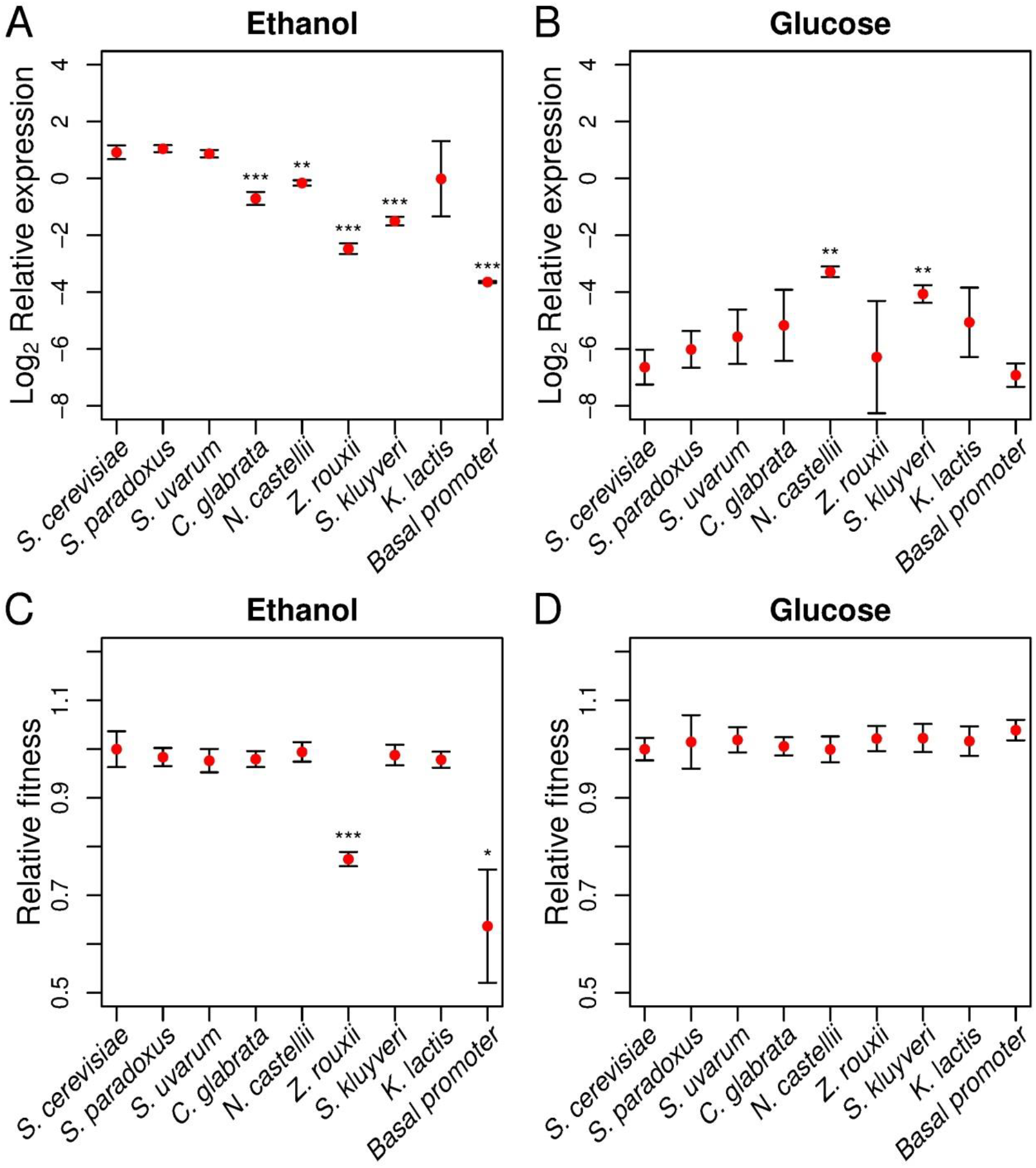
Gene expression and fitness of *MLS1* promoters in steady state conditions. Relative expression shows expression of each species’ *MLS1* promoter and *S. cerevisiae*’s core promoter relative to the housekeeping gene *ACT1* on a log_2_ scale in (A) 3% ethanol and (B) 2% glucose. Relative fitness represents the growth rate of each strain relative to a reference competitor strain in (C) 3% ethanol and (D) 2% glucose. Error bars indicate one standard deviation, significant differences in comparison to *S. cerevisiae* are shown for Bonferroni corrected P<0.05 (*), P<0.01 (**) and P<0.001 (***).

To gauge the extent to which the *S. cerevisiae MLS1* promoter is influenced by known binding sites, we compared the *S. cerevisiae* promoter to: 1) a promoter lacking the *MLS1* proximal Cat8/Sip4 site, 2) a promoter lacking both Cat8/Sip4 sites, 3) a promoter lacking the Mig1 site, and 4) a core promoter containing only the proximal 186 bp of promoter, which includes TATA but lacks the Abf1, Cat8 and Mig1 sites (Figure S2). While deletion of either the proximal or both Cat8/Sip4 sites did not affect expression (Figure S3A), the core promoter was expressed at much lower levels in ethanol (Figure 2A). However, the core promoter still caused a 9.7-fold increase in expression in ethanol compared to glucose, similar to the level of activation found for the promoters of *N. castellii, L. kluyveri* and *Z. rouxii* (Figure 2A-B; Table S2). Similar to a previous study (Caspary 1997), the Mig1 deletion caused a loss of repression (Figure S3B).

### Fitness is maintained despite changes in gene expression levels

We tested whether any of the differences in expression affect fitness by competing each strain bearing a different *MLS1* promoter with a common reference strain in either glucose or ethanol. There were no significant differences in fitness between the *S. cerevisiae* promoter and that of any other species except for *Z. rouxii* in ethanol (Figure 2C and 2D). This indicates that in relationship to gene expression levels in ethanol, there is a fitness plateau and a sharp cliff between the expression levels of *L. kluyveri* and *Z. rouxii* (Figure 2A). The fitness of the promoter deletions are largely consistent with their measured expression levels (Figure 2 and S3). Deletion of all binding sites except the core promoter had a large impact on fitness, consistent with its low expression level, and deletion of the Sip4/Cat8 and Mig1 sites had little to no impact on fitness in ethanol and a slight increase in fitness in glucose (Figure S3C). Compared to an *S. cerevisiae* strain with *MLS1* at its endogenous locus, the transgenic *S. cerevisiae* allele of *MLS1* at the *URA3* locus exhibited a slight decrease in expression in ethanol, a slight increase in fitness in ethanol, and a slight decrease in fitness in glucose (Figure S3).

### Dynamic expression and fitness in fluctuating environments

Our previous measurements of gene expression levels and fitness were done during steady-state growth after cells were allowed to condition themselves to growth on glucose or ethanol. However, the dynamic response of a promoter to an altered environment may be as important to fitness as expression levels under steady-state conditions. To examine the temporal dynamics of each species’ promoter we measured expression following a switch from growth on glucose to ethanol. Similar to steady state levels, expression dynamics are conserved within the *Saccharomyces* species as is apparent from the consistent response over time (Figure 3A). For all the non-*Saccharomyces* species we observed a smaller increase in expression between 0 and 15 minutes after switching to growth on ethanol (Figure 3B-C; Table S3). *Z. rouxii* and *L. kluyveri* also showed a smaller increase in expression between 15 and 30 minutes (Figure 3B-C; Table S3). While the dampened response of non-*Saccharomyces* species to ethanol is consistent with their lower steady-state levels, the absolute expression level at 15 minutes was only different between *S. cerevisiae* and *C. Glabrata* (Table S4).

**Figure 3.**
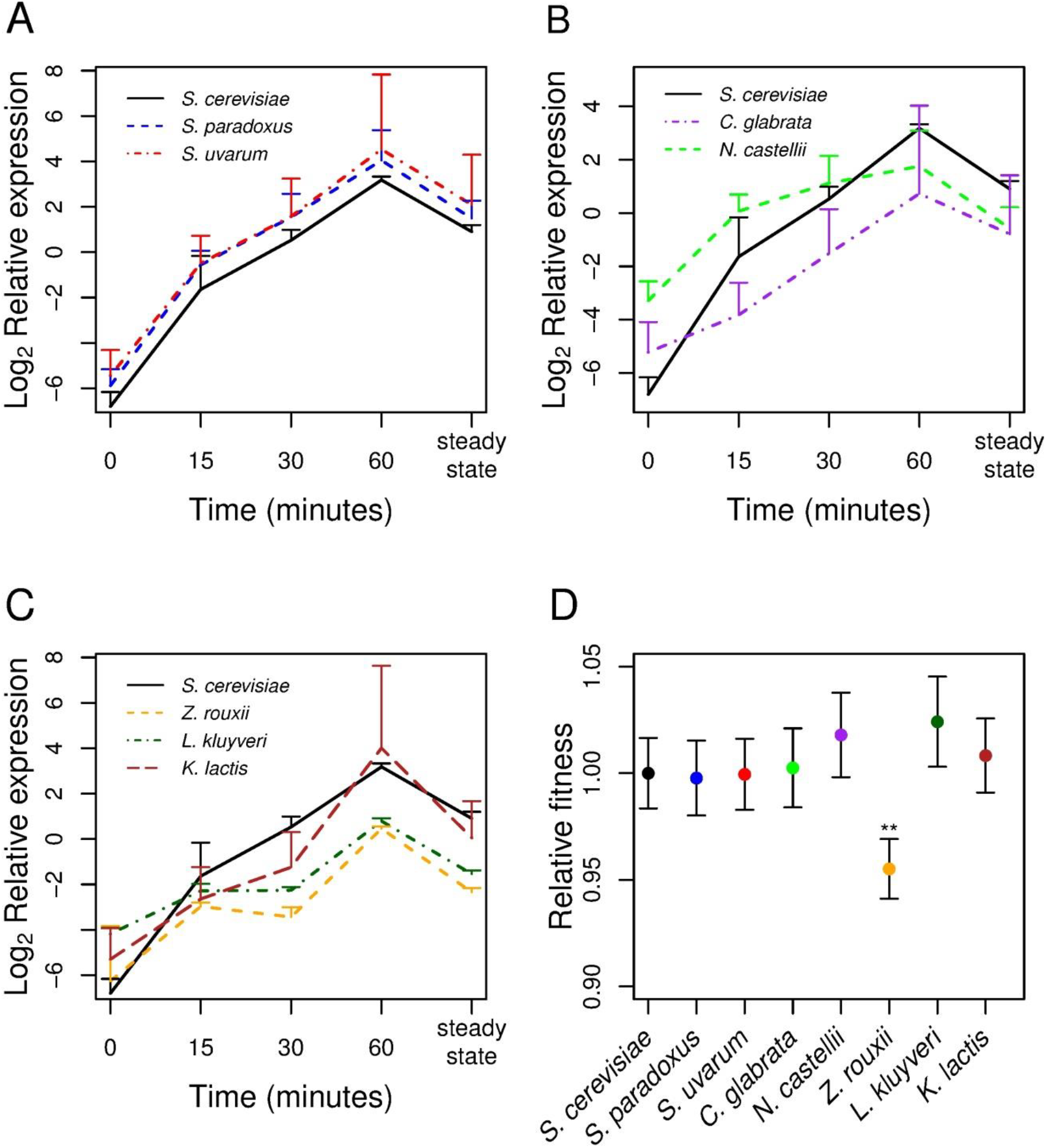
Gene expression and fitness of *MLS1* promoters in a fluctuating environment. Changes in *MLS1* expression for the eight species promoter constructs are shown in A-C and are divided into (A) *Saccharomyces* species, (B) post-whole genome duplication species, and (C) pre-whole genome duplication species. Relative expression levels represents the level of *MLS1* relative to the housekeeping gene *ACT1* on a log_2_ scale. Time point 0 is expression in complete media with 2% glucose and subsequent time points are expression levels 15, 30, and 60 minutes after being placed in complete media with 3% ethanol. The steady state time point indicates expression after growing for over 24 hours in ethanol media. Error bars indicate one standard deviation. Relative fitness of each *MLS1* species promoter construct (D) measured as the growth rate of each strain relative to a fluorescent competitor strain after three days of sequential competition in complete media with 3% ethanol plus 0.2% glucose. Bonferroni corrected p-values are indicated for P<0.05 (*), P<0.01 (**) and P<0.001 (***).

Given the different expression dynamics, we tested whether the *MLS1* promoters cause fitness differences in an environment fluctuating between glucose and ethanol. Only *Z. rouxi* showed significantly reduced fitness (Bonferroni corrected p-value < 0.05, Figure 3D), the only species with lower fitness in steady state ethanol (Figure 1C). The higher fitness of *Z. rouxii* in a fluctuating compared to constant environment is likely a result of the competition including growth in glucose where no fitness defect was measured. In the fluctuating environment there was a slightly higher fitness for *L. kluyveri*, potentially caused by the weaker *MLS1* repression in glucose enabling *L. kluyveri* to start growing earlier. The other species with a significant loss of repression in glucose, *N. castellii* (Figure 1B), had higher fitness in the fluctuating environment that is close to significant.

## DISCUSSION

Knowing how gene expression affects fitness is important to interpreting patterns of gene expression divergence. Using *MLS1* promoters from eight yeast species, we find that large differences in gene expression levels do not generate detectable fitness effects. However, we also find a large drop in fitness below a certain low level of expression, implying that the *S. cerevisiae MLS1* promoter resides on a fitness plateau. The high fitness of various configurations of binding sites present in different species provides further experimental support for the flexibility of the *cis*-regulatory code.

### Conservation of carbon source response combined with divergence in expression levels

Similar to previous promoter studies (Gordon and Ruvinsky 2012), we find *MLS1* promoters are conserved in their ability to respond to glucose and ethanol but also exhibit loss of both activation and repression as divergence between these elements and *S. cerevisiae* increases. Also consistent with a previous study of interspecific divergence (Tirosh *et al.* 2008), we find little correspondence between changes in binding sites and expression levels. First, deletion of one or both Cat8/Sip4 sites did not affect expression and there is no strong correlation between the summed binding affinity of a promoter and its expression level (Table S1). A previous study (Caspary *et al.* 1997) found that mutation of the proximal Cat8/Sip4 site or both Cat8/Sip4 sites caused a 28% and 80% reduction in expression, respectively. One explanation for for why we did not find effects for these sites is that our deletion constructs altered the spacing of other binding sites, e.g. Abf1 sites were brought close to TATA. However, the different results could also be a consequence of Caspary *et al*. (1997) measuring expression from a high copy episomal plasmid whereas we measured expression from a construct integrated into the *URA3* locus. A second line of evidence for the importance of sequences besides Cat8/Sip4 sites is that the core promoter still yielded a 9.7-fold increase in expression in ethanol compared to glucose (Table S2). However, we did find effects associated with the Mig1 binding site: the Mig1 deletion caused a loss of repression in the *S. cerevisiae* promoter and *N. castellii* had the highest expression in glucose and also lacked a Mig1 site.

### **The fitness-expression function in *S. cerevisiae***

Previous work has shown conditional fitness costs and benefits of Lac expression in bacteria (Dekel and Alon 2005; Perfeito *et al.* 2011), and a fitness plateau for *LCB2* expression in yeast (Rest *et al*. 2013). Results for *MLS1* expression differ from *LCB2* in that endogenous *LCB2* expression levels occurred at the edge of the fitness cliff whereas no detectable loss of fitness occurred for up to a 5.4-fold (*L. kluyveri*) drop below wildtype levels for *MLS1*. We put forth three explanations for the high level of *MLS1* expression in *S. cerevisiae*. First, low *MLS1* expression levels may cause reduced fitness in conditions other than those measured. For example, *MLS1* is required for sporulation and low expression could reduce or alter sporulation efficiency. Second, high *MLS1* expression could be maintained by small fitness effects that are not detectable by our assays. Because purifying selection can occur on selection coefficients as small as the inverse of the effective population size, the fitness plateau could be covered with undetectable hills. In support of this possibility, there is good evidence for purifying selection on *MLS1* binding sites within the *Saccharomyces* species (Figure S4; Doniger and Fay 2007). Third, the fitness-expression function may only have a small plateau in other genetic backgrounds. Strain differences in *LCB2* expression imply that genetic background modulates the fitness-expression function (Rest *et al.* 2013).

What types of expression changes affect fitness? In addition to steady-state levels, prior work in yeast showed that noise in *TDH3* expression affects fitness (Metzger *et al.* 2015). *MLS1* like other TATA-containing promoters is characterized by large variation in cell-to-cell levels of expression, which may provide a fitness advantage under fluctuating environments through bet hedging (Thattai *et al.* 2004; Kaern *et al.* 2005). However, we found fitness effects in a fluctuating environment to mirror and be smaller than those under steady-state conditions.

One limitation of our approach is that we used heterologous expression and fitness assays. As such, it is possible that *MLS1* promoters from the distantly related yeast species do not have reduced activation and repression in their endogenous genome. Both the extensive *cis*-*trans* expression interactions found to occur between species (McManus *et al.* 2010; Swain Lenz *et al.* 2014) and the dependency of the fitness-expression function on strain background (Rest *et al*. 2013) indicate that endogenous *MLS1* expression in other species may not be the same as that measured in *S. cerevisiae*. However, the expression levels, fitness effects and configuration of binding sites are directly relevant to the strain of *S. cerevisiae* in which it was measured and interpreting what changes in the *MLS1* promoter are likely to be tolerated.

In conclusion, our finding of a fitness plateau for *MLS1* expression provides an explanation for divergence in gene expression levels and configurations of binding sites without an overall change in carbon source response. While it is unknown whether most genes have a fitness-expression plateau, the paucity of haploinsufficient genes in yeast (3%) implies most genes can tolerate a 2-fold change in expression without major fitness defects (Deutschbauer *et al.* 2005). Current models for the evolution of *cis*-regulatory sequences, including binding site turnover (Ludwig *et al.* 2000) and rewiring of transcription factors (Perez and Groisman 2009), suppose that the diversity of orthologous *cis-* regulatory sequences is a product of compensatory changes. However, when fitness effects are small or absent, many changes in *cis*-regulatory sequences may evolve under a nearly neutral model despite their effects on gene expression.

## MATERIALS AND METHODS

Binding site and nucleosome predictions – Position weight matrices (PWM) for Abf1, Sip4 and Mig1 were obtained from MacIsaac *et al. (*2006). The PWM for Cat8 was obtained from a curated list of motifs (Soontorngun *et al* 2007), and the PWM for TATA (NHP6A) was from Zhu *et al*. (2009). Sequences were searched for binding sites using Patser (Hertz and Stormo 1999). Only binding site scores below a ln(p-value) of 7 were considered, where the p-value is the expected probability of a random match to the binding site (Hertz and Stormo 1999). Nucleosome occupancy probability was predicted for each *MLS1* promoter (Kaplan *et al.* 2009). The temperature and histone concentration parameters were set to 1 and 0.03, respectively, as in Kaplan *et al.* (2009).

Species promoter constructs - *MLS1* promoter regions from 8 species were placed into an *S. cerevisiae* background. First, the pRS306-ScMLS1 plasmid was constructed by inserting *MLS1* from *S. cerevisiae* (S288c) into the integrative plasmid pRS306 (Sikorski and Hieter, 1989). The *MLS1* region from S288c includes the 893 bp noncoding region upstream of *MLS1* as well as the 305 bp region downstream of the *MLS1* translation stop site. Second, the promoter of *S. cerevisiae MLS1* in pRS306-ScMLS1 was then replaced by the *MLS1* promoter in seven other yeast species in the following manner. *MLS1* promoter regions were defined as the noncoding region upstream of the *MLS1* start codon to the beginning of the next coding region. In the cases of *S. castellii* and *Z. rouxii*, the predicted intergenic regions were short (575 bp for *Z. rouxii* and 250 bp for *S. castellii*) and therefore, the region used for these two promoters was ∼ 1 kb upstream of the *MLS1* start codon. The promoter region of *MLS1* from each species (Table S5) was PCR amplified (see File S1 for primers) and subcloned into the pRS306-ScMLS1 plasmid using the Gibson Assembly method (New England Biolabs, Ipswich, MA). The promoter region as well as the *S. cerevisiae MLS1* coding region were sequence confirmed for each construct.

Binding site deletions – Binding sites were deleted by removing the region surrounding each binding site from the promoter. Regions deleted are shown in Figure S2. Deletions were generated by amplifying the pRS306-ScMLS1 plasmid with segments of the promoter missing. Primers contained BglII sites on their 5’ end (see File S1). After amplification, the PCR product was digested with BglII and ligated back together to form a circular plasmid. The *S. cerevisiae MLS1* promoter deletions and coding region were sequence confirmed.

Plasmid integrations - The endogenous *MLS1* coding region from the strain YJF186 (YPS163 oak isolate, Mat *a*, HO::dsdAMX4, ura3-140) was deleted by replacement with the KANMX4 cassette to generate the strain YJF604. All pRS306 based plasmids described above were cut in the URA3 coding region with StuI and integrated into YJF604 using lithium acetate transformation (Geitz and Woods 2002) and selected on plates lacking uracil. The competitor strain containing yellow fluorescent protein (YFP) was generated by integrating a YFP-NATMX4 plasmid containing homology to the HO locus (received from R. Kishony) into YJF186.

Competitive fitness assays – Fitness was estimated by competing each strain against a YFP marked reference strain. For each competition six biological replicates (independent transformants) of each integrated construct were competed against the YFP competitor at 30°C at 300 rpm in 3 mL media in 18 x 150 mm glass tubes. Ethanol (3%), glucose (2%) and mixed carbon source (3% ethanol 0.2% glucose) competitions were carried out in complete medium (CM: 0.67% (w/v) nitrogen base with ammonium sulfate and amino acids) with the specified carbon sources. All strains were acclimated to each growth medium prior to competition by 3 days of growth, with cells resuspended in fresh medium after each day. The YFP competitor strain was mixed with each culture at a 50:50 ratio at a starting cell density of OD_600_ = 0.07. Competitions in CM(3% ethanol) and CM(3% ethanol + 0.2% glucose) were carried out for 2 days with resuspension in fresh medium after every 23hrs of competition. Competitions in CM(2% glucose) were carried out for 1 day with resuspension in fresh medium after every 11hrs of competition. The proportion of YFP positive strains was determine at the beginning and end of each competition. Cells from each culture were also diluted to an OD_600_=0.2 in sheath fluid and run on a Beckman Coulter FC 500 MPL flow cytometer (Beckman Coulter, Brea, CA) to distinguish between fluorescent and non-fluorescent cells.

Fitness calculations *–* Fitness measurements were calculated using w_i_= (*N*_1_/*Y*_1_) − *ln(N*_0_/*Y*_o_) as in Hartl and Clark (1997), where *Y*_0_ and *N*_o_are the starting frequencies of the YFP strain and the non fluorescent competitor strain, respectively. Here, *N*_0_ = 1 − *Y*_0_. Similarly, *N*_1_and *Y*_1_represent these frequencies at the end of the competition. Relative fitness of a given strain *i* is equal to 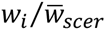, where 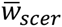 is the average fitness of the *S. cerevisiae* strains.

MLS1 mRNA expression analysis − *MLS1* measurements in steady-state growth were measured as follows. Using four of the same replicates for each promoter from the competition, each strain was acclimated and cells were sampled after 4 hrs in fresh CM(3% ethanol) or CM(2% glucose) medium (OD_600_ = 1) on the third day. The equivalent of 1 mL cells at an OD600 of 0.3 were sampled. Cells were centrifuged, supernatant was removed, pellets were frozen in liquid nitrogen, and stored at -80°C.

*MLS1* mRNA expression during the switch from glucose to ethanol was obtained from 4 time-points. After 3 days acclimation cells were placed in 3 mL CM(2% glucose) at an OD_600_=1 and grown for 4 hours. Cells were centrifuged for 30 seconds at 3,000 rpm, supernatant was removed and cells were washed with 1 mL CM(3%ethanol) and centrifuged again. Supernatant was removed and cells were resuspended in 3 mL of CM(3% ethanol) and cultures were placed in the incubator. Cells were then sampled 15, 30, and 60 minutes after cells were initially placed into CM (3% ethanol), centrifuged, and pellets were frozen in liquid nitrogen and stored at -80C.

*MLS1* expression was measured using QuantiGene (Affymetrix, Santa Clara, CA) following manufacturers instructions. 200 μL Homogenization buffer (Affymetrix) was added to each pellet, resuspended, centrigued, and supernatant was removed. Pellels were resuspended in 100 μL of ZYM buffer (Clontech, Mountain View, CA) and 10 μL zymolase (Clontech) and allowed to digest for 1 hour at 30°C at 300 rpm. After digestion, 150 μL of Homogeniztion buffer was added to each well. The content of each well was then diluted 1:100 in homogenization buffer. Next, 40 μL of these 1:100 diluted samples were added to 60 μL of ‘working bead mix’ described in steps 4-6 of the ‘Purified RNA or in vitro Transcribed RNA’ protocol in the QuantiGene 2.0 Plex Assay User Manual (Panomics Solutions P/N 16659 Rev.C 020912). The ‘Purified RNA or in vitro Transcribed RNA’ protocol was then followed exactly from step 7 onwards. Probes were designed to the *MLS1* and *ACT1* coding regions of *S. cerevisiae*. 40 μL of the 1:100 diluted samples was added to 60 μL of mastermix. Measurements were obtained on a Bio-Plex 200 System (Life technologies, Carlsbad, CA) and analyzed using the Bio-Plex Manager™ 6.1 software. Standard curves for each analyte were generated by a 4-fold serial dilution of one of the *S. cerevisiae MLS1* promoter strains sampled in ethanol media.

Statistical analysis of fitness and expression – Six biological replicates (independent integrations at the URA3 locus) of each promoter construct were measured for fitness. Four biological replicates of each promoter were measured for expression in steady state glucose and ethanol. Outliers from each group were removed using the Grubbs’ test (P-value <0.05). Significant differences were measured by t-tests with unequal variance. Bonferonni correction was used for the seven hypotheses that another species promoter was different than the *S. cerevisiae* promoter. For measurements of the dynamics of gene expression from glucose to ethanol, three biological replicates were used and no outliers were removed. A nested ANOVA was used to measure the differences between each species’ promoter at each time point as well as the rate of change (slope) between each time point. This was done in R where level∼(species/time).

## ACKNOWLEDGEMENTS

The authors would like to thank Xueying Li, Ping Liu, Kim Lorenz, Linda Riles, Ching-Hua Shih, and Katie Williams with help on competition assays and Holly Brown with help on the QuantiGene assay. The research was supported by an NIH training grant (GM007067) to ACB and GM08669 to JCF.

